# The PRRA insert at the S1/S2 site modulates cellular tropism of SARS-CoV-2 and ACE2 usage by the closely related Bat raTG13

**DOI:** 10.1101/2020.07.20.213280

**Authors:** Shufeng Liu, Prabhuanand Selvaraj, Christopher Z. Lien, Wells W. Wu, Chao-Kai Chou, Tony T. Wang

## Abstract

Biochemical and structural analyses suggest that SARS-CoV-2 is well-adapted to infecting human and the presence of four residues (PRRA) at the S1/S2 site within the Spike protein may lead to unexpected tissue or host tropism. Here we report that SARS-CoV-2 efficiently utilized ACE2 of 9 species except mouse to infect 293T cells. Similarly, pseudoviruses bearing spike protein derived from either the bat raTG13 or pangolin GX, two closely related animal coronaviruses, utilized ACE2 of a diverse range of animal species to gain entry. Removal of PRRA from SARS-CoV-2 Spike displayed distinct effects on pseudoviral entry into different cell types. Strikingly, insertion of PRRA into the raTG13 Spike selectively abrogated the usage of horseshoe bat and pangolin ACE2 but conferred usage of mouse ACE2 by the relevant pseudovirus to enter cells. Together, our findings identified a previously unrecognized effect of the PRRA insert on SARS-CoV-2 and raTG13 spike proteins.

## Introduction

The outbreak of the severe acute respiratory syndrome coronavirus 2 (SARS-CoV-2 or 2019-nCoV) has quickly turned into a global pandemic. Multiple studies have suggested a zoonotic origin of SARS-CoV-2, highlighting the potential of coronaviruses to continuously evolve in wild animal reservoirs and jump to new species (Andersen et al., 2020; Zhou et al., 2020a). Various bat species are believed to be reservoirs of progenitors of diverse coronaviruses (CoVs) including the severe acute respiratory syndrome (SARS)-CoV (SARS-CoV) and SARS-CoV-2. Indeed, the closest virus to SARS-CoV-2 found up to date is the bat raTG13 with over 96% identity at genomic level (Zhou et al., 2020b). A newly reported bat RmYN02 shares 93.3% nucleotide identity with SARS-CoV-2 genomewide and 97.2% identity in the 1ab gene (Zhou et al., 2020a). Besides bat CoVs, pangolin/GD/2019 CoV and pangolin/GX/P5L/2017 CoV share up to 90% and 85.2% sequence identity with SARS-CoV-2, respectively (Liu et al., 2020; Zhang et al., 2020). While the origin of the novel coronavirus appears to be in bat reservoirs, there is still no definitive evidence of an intermediate host that could have transmitted the virus to humans.

The coronavirus spike protein plays a pivotal role in mediating coronavirus entry (Belouzard et al., 2012; Li, 2016). The ectodomain of SARS-CoV spike protein is divided into two subunits, S1 and S2. SARS-CoV S1 contains a receptor-binding domain (RBD) that specifically binds to human angiotensin-converting enzyme 2 (ACE2) whereas S2 subunit contains the fusion peptide (Li et al., 2005; Li et al., 2003). Like SARS-CoV, the SARS-CoV-2 spike protein in pre-fusion conformation is a homotrimer, with three S1 heads sitting on top of a trimeric S2 stalk (Shang et al., 2020b; Walls et al., 2020; Wrapp et al., 2020). Based on a structure of the SARS-CoV-2 spike ectodomain containing two stabilizing mutations, it was observed that at a given time, only one of three RBD is in an upward position for receptor binding (Wrapp et al., 2020). However, a very recent structural study of a full-length, fully wild-type form of the SARS-CoV-2 S protein suggested that the prefusion structure is in the “all-down” configuration (Yongfei Cai, 2020). Interaction between RBD and ACE2 is thought to lock the SARS-CoV-2 spike protein in a conformation that makes it susceptible to multiple cellular proteases (Yan et al., 2020), including cell surface protease TMPRSS2 and lysosomal proteases cathepsins (Hoffmann et al., 2020; Millet and Whittaker, 2015). Proteolytic cleavage of SARS-CoV spike at the S1/S2 boundary causes S1 to dissociate so that S2 undergoes a dramatic structural change to induce fusion(Millet and Whittaker, 2015). All this evidence indicates that SARS-CoV-2 may follow the footstep of SARS-CoV to enter host cells.

In comparison to SARS-CoV and other betacoronaviruses, the spike protein of SARS-CoV-2 displays two distinct features: 1) SARS-CoV-2 appears to have been well adapted for binding to the human receptor ACE2, despite significant sequence divergence in its RBD region in comparison to that of SARS-CoV (Lan et al., 2020; Wu et al., 2020). 2) A distinct four amino acid insert within the spike (S) protein (underlined, SPRRAR↓S) between the S1 receptor binding subunit and the S2 fusion subunit presumably creates a functional polybasic (furin) cleavage site. This novel S1/S2 site of SARS-CoV-2 Spike is hypothesized to affect the stability, transmission and/or host range of the virus (Jaimes et al., 2020a; Jaimes et al., 2020b).

In this study, we set out to determine 1) if SARS-CoV-2 infects cells via diverse animal ACE2; 2) how the PRRA insert at the S1/S2 site affects the cellular and host range of SARS-CoV-2 and its closely related RaTG13. Our findings not only demonstrated the wide usage of ACE2 from diverse animal species by SARS-CoV-2, raTG13 and pangolin GX, but also revealed a striking effect of insertion of PRRA on raTG13 spike protein regarding its usage of ACE2 derived from horseshoe bat, pangolin and mouse.

## Results

### Efficient infection of cells expressing ACE2 of diverse species by SARS-CoV-2

Structural and biochemical studies have suggested that SARS-CoV-2 may have been well adapted in human before the outbreak in Wuhan, China (Andersen et al., 2020; Shing Hei Zhan, 2020). Despite significant variation in the receptor binding motif (RBM) from that of the SARS-CoV spike, SARS-CoV-2 spike protein appears to have evolved a mechanism to efficiently utilize human ACE2 to gain entry. Such a feature had not previously been recognized. Besides human, SARS-CoV-2 reportedly infects domesticated pets, such cats and dogs (Shi et al., 2020), and wild animals, such as mink (Nadia Oreshkova, 2020), raising the concern of cross species transmissions. To test whether SARS-CoV-2 may use ACE2 of different species to infect cells, we transfected 293T cells to express ACE2 from 10 different species, including human, mouse, horseshoe bat, pangolin, Syrian (SYR) hamster, Chinese (CHN) hamster, African green monkey (Agm), bovine, pig, ferret. Cells were subsequently infected by SARS-CoV-2 that expresses the monomeric neon green fluorescent protein (mNeonGreen) (Xie et al., 2020). Except mouse ACE2, expression of all ACE2 (Fig. S1A) rendered cells susceptible to SARS-CoV-2 infection (Fig. 1).

**Figure 1.**
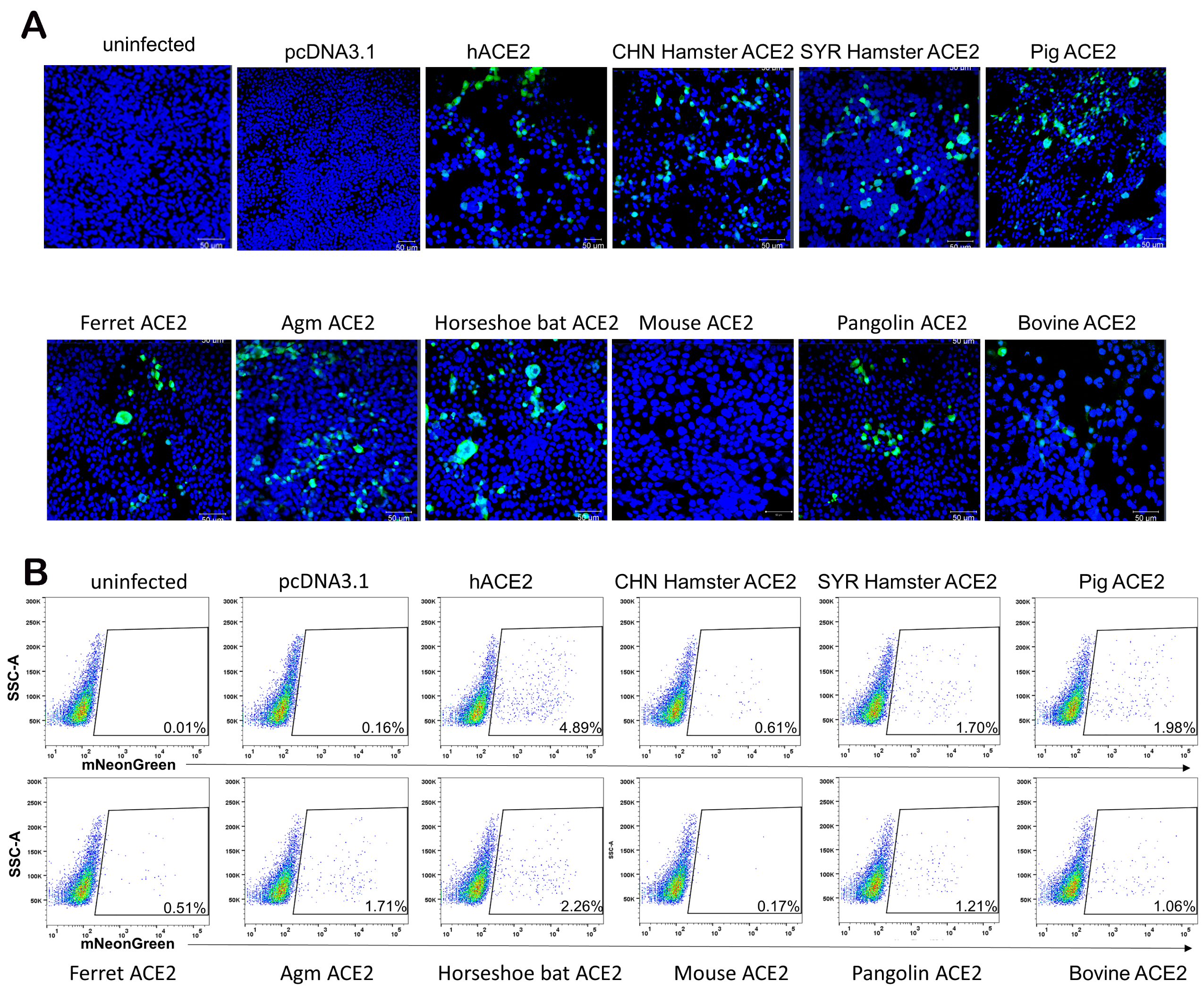
ACE2 orthologs of diverse species mediate SARS-CoV-2 infection. (A) 293T cells transfected with plasmids expressing 10 ACE2 orthologs or empty vector and then infected by icSARS-CoV-2-mNG at multiplicity of infection of 1 for 48 hours. Images were captured by a Zeiss LSM 880 laser scanning microscope. The scale bar = 50 μm. (B) Quantification of infection by flow cytometry. Cells from (A) were fixed in 4% paraformaldehyde and then mNG positive cells were quantified by flow cytometric analysis.

### PRRA-led proteolytic cleavage of SARS-CV-2 Spike protein

Next, we sought to investigate the role of the PRRA insert in SARS-CoV-2 entry. To confirm cleavage by cellular proteases at this site, we transfected 293T cells with constructs expressing the wild type SARS-CoV-2 spike (SARS-CoV-2 S), a mutant with PRRA removed (SARS-CoV-2 S ΔPRRA), the Bat raTG13 spike (raTG13 S), an insertion mutant containing the PRRA (raTG13 S+PRRA), and the Pangolin GX Spike protein (Pangolin GX S) (Fig. 2A). We used a mouse monoclonal antibody that recognizes the S2 domain of both SARS-CoV spike and SARS-CoV-2 spike proteins for detection. A partial cleavage was noticed when SARS-CoV-2 Spike was expressed in 293T cells and the deletion of PRRA abolished the cleaved form (Fig. 2B).

**Figure 2.**
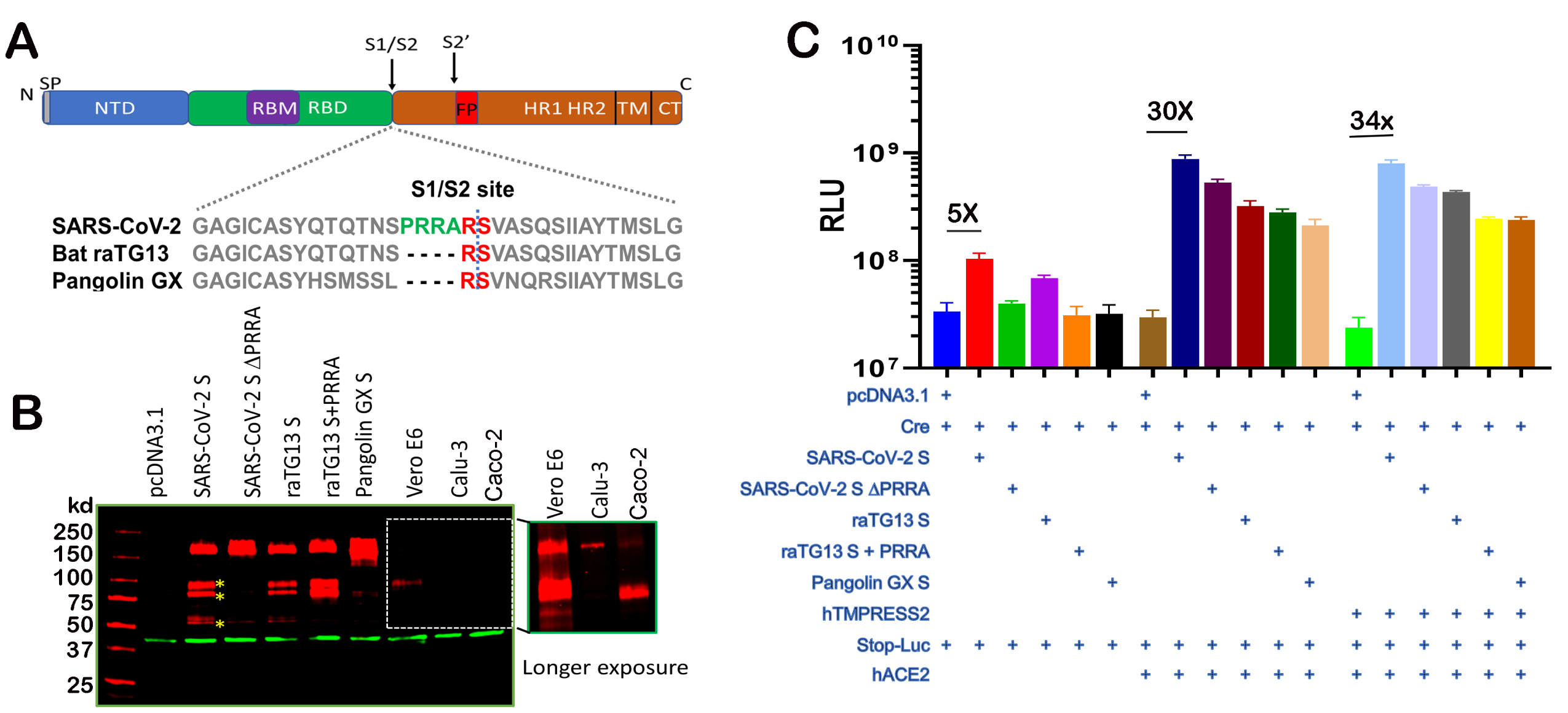
PRRA-led proteolytic cleavage of SARS-CV-2 Spike protein and effect on fusion. (A) Organization of SARS-CoV-2 spike protein and the sequence alignment at S1/S2 site with raTG13 and Pangolin GX S. (B) 293T cells transfected with SARS-CoV-2 S, SARS-CoV-2 S ΔPRRA, the Bat raTG13 spike (raTG13 S), an insertion mutant containing the PRRA (raTG13 S+PRRA), and the Pangolin GX Spike protein (Pangolin GX S), lane 3-7. SARS-CoV-2 S infected Vero E6, Calu-3 and Caco-2, lane 7-10. A longer exposure of this part of the gel is included. Red, anti-S antibody; green, anti-β-actin antibody. (C) Cell–cell fusion mediated by CoVs S protein. 293T cells expressing Stop-Luc and/or ACE2, ACE2/TMPRSS2 (acceptor cells) were mixed at a 1:1 ratio with donor cells expressing Cre, CoV S, or both to initiate cell-cell fusion. Data are represented as mean ± SEM.

Interestingly, raTG13 spike was partially cleaved even in the absence of PRRA. Insertion of PRRA insertion into raT13 S further enhanced the cleavage by about 2-fold. Lastly, pangolin GX spike protein did not show similar cleavage pattern in transfected 293T cells. To confirm that the cleavage occurs in the context of infection, western blotting was performed using SARS-CoV-2 infected VeroE6 cell lysate, similar cleavage pattern was observed (Fig. 2B). Of note, the extent of the observed cleavage appears to vary significantly among susceptible cell lines. While the ratio of the cleaved over noncleaved Spike appears to be comparable between 293T and Vero E6, infected human lung epithelial cell line Calu-3 and intestinal cell line Caco-2 by SARS-CoV-2 yielded banding patterns that indicate a rather complete cleavage of the spike protein in Caco-2 and nearly no cleavage in Calu-3 cells (Fig. 2B and Fig. S1B).

A pre-cleaved spike protein at S1/S2 could confer unexpected fusogenic capability on the plasma membrane and allow coronaviruses undergo receptor-independent entry (virus–cell fusion) (Li, 2016; Miura et al., 2008). To assess the Spike protein-mediated membrane fusion, we set up a quantitative cell-cell fusion assay in which 293T acceptor cells, containing a loxP-flanked STOP cassette that blocks transcription of the downstream luciferase reporter gene, were cocultured with Cre-expressing donor cells. In this assay, fusion between donor and acceptor cell membranes removes the STOP cassette and hence permits luciferase production. Various spike constructs were transfected into donor cells and then the hACE2 plasmid was transfected into the acceptor cells. Notably, even in the absence of hACE2, SARS-CoV-2 spike induced about 5-fold over the background (Fig. 2C). Deletion of PRRA from SARS-CoV-2 spike completely abolished this basal level activity. In the presence of ACE2, all spike proteins induced cell-cell fusion to varying extents (up to 30-fold) above background with the wild type SARS-CoV-2 Spike being the most efficient one. Surprisingly, insertion of the novel PRRA into raTG13 S and deletion of PRRA from SARS-CoV-2 S both reduced fusogenic ability in the presence of ACE2 in comparison to their wild type counterparts. Finally, addition of TMPRSS2 only marginally affected the Spike-driven cell-cell fusion (Fig. 2C).

### The PRRA insert at the S1/S2 site distinctly modulates cell susceptibility to SARS-CoV-2 pseudovirions

Next, we sought to investigate the impact of PRRA insert on virus entry. To this end, we packaged pseudoviruses bearing SARS-CoV-2 S, SARS-CoV-2 S ΔPRRA, raTG13 S, raTG13 S+PRRA, and Pangolin GX S. Again, we noticed a basal level of infection by pseudovirus bearing SARS-CoV-2 Spike into 293T cells even in the absence of ACE2 (Fig. S2A). This finding is consistent with the observed basal level of cell-cell fusion that is mediated by SARS-CoV-2 Spike. Surprisingly, pseudovirus carrying SARS-CoV-2 S ΔPRRA consistently infected Vero E6 and 293T-hACE2 cells at about half to one log10 higher than the wild type SARS-CoV-2 spike (Fig. 3A and Fig. S2B). By contrast, infection of the human lung epithelial cell line Calu-3 as well as the human intestinal cell line Caco-2 by the same pseudoviruses were reduced by 90-50%, respectively, compared to pseudoviruses bearing the wild type SARS-CoV-2 S or raTG13 S (Fig. 3B&C). Among the five pseudoviruses, bat raTG13 spike appears to be the least efficient one in mediating pseudovirus entry through hACE2. Insertion of PRRA into raTG13 Spike further reduced the pseudovirus entry into all cell types that were tested (Fig. 3A-C). Pseudovirus bearing the Pangolin spike is highly infectious on 293T-hACE2 (Fig. 3A). Interestingly, the spike protein displayed on pseudoviruses bearing SARS-CoV-2 S, raTG13 S, raTG13 S+PRRA were almost completely cleaved, whereas it was noncleaved on pseudovirions carrying the SARS-CoV-2 S ΔPRRA (Fig. 3D). Pangolin GX S on the pseudovirions appears to display an entirely different pattern with potentially partial cleavage (Fig. 3D). Lastly, expression of TMPRSS2 increased the infectivity of pseudoviruses bearing SARS-CoV-2, raTG13 S, and raTG13 S+PRRA by about one log, but not those bearing SARS-CoV-2 S ΔPRRA and the Pangolin GX S (Fig. 3E).

**Figure 3.**
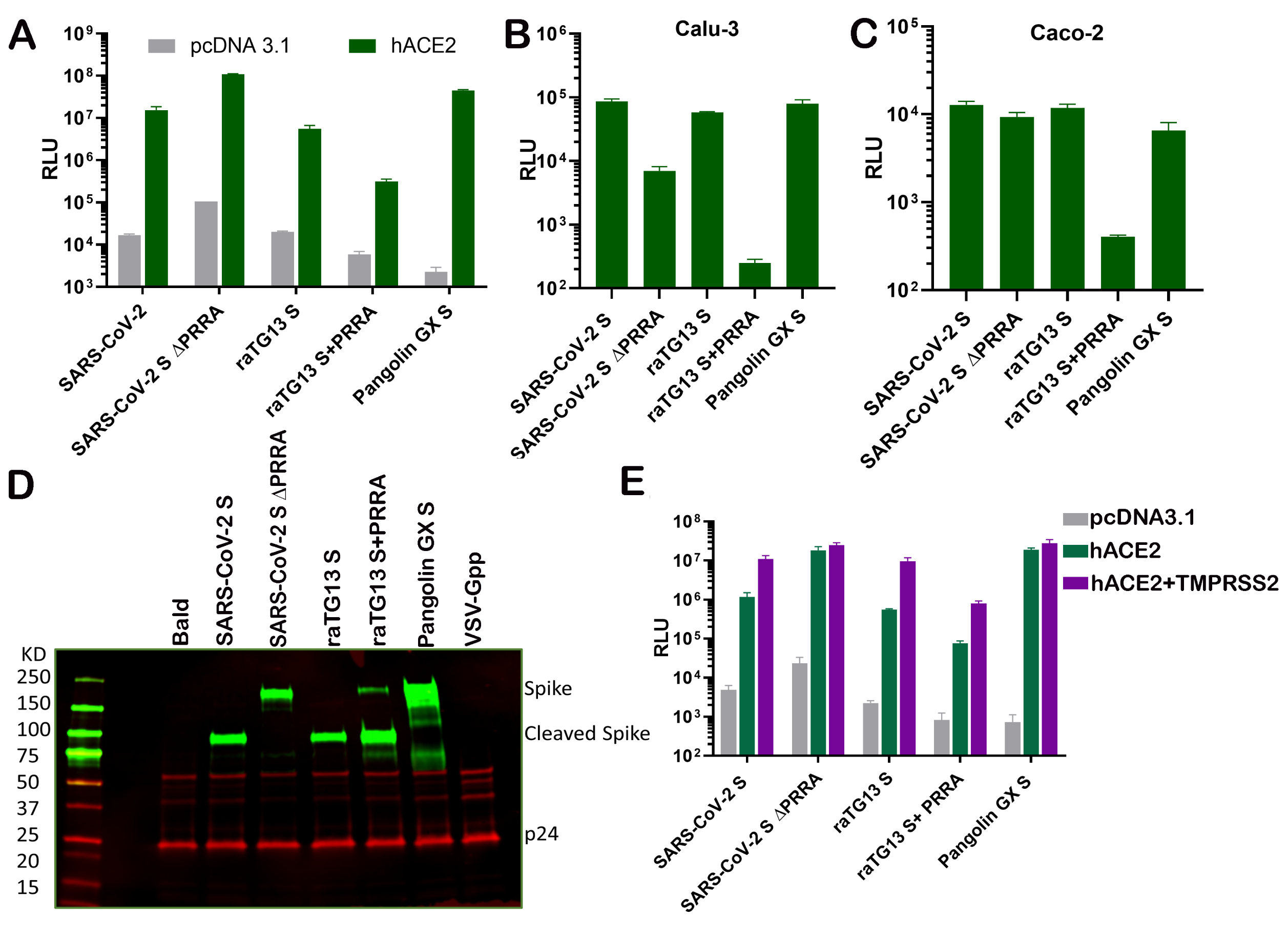
The PRRA insert at the S1/S2 site distinctly modulates cell susceptibility to SARS-CoV-2 pseudovirions. (A) Entry of HIV pseudotyped with SARS-CoV-2 S, SARS-CoV-2 S ΔPRRA, bat raTG13 S, raTG13 S+PRRA and Pangolin GX S into 293T cells transiently expressing ACE2 (A), Calu-3 (B) and Caco-2 (C). (D) Western blot analysis of pseudovirions bearing SARS-CoV-2 S, SARS-CoV-2 S ΔPRRA, bat raTG13 S, raTG13 S+PRRA and Pangolin GX S. Red, anti-p24 antibody; green, anti-Spike antibody. (E) Infection of pseudovirons harboring the indicated viral glycoproteins into 293T cells transfected with ACE2 alone or in combination with TMPRSS2. Data are represented as mean ± SEM.

### The novel S1/S2 site alters the dependence of raTG13 pseudovirus on ACE2 of different species

The PRRA insert between S1/S2 boundary of SARS-CoV-2 Spike is suspected to alter stability, transmission and even host range of SARS-CoV-2. Furthermore, recent studies suggest that a bat virus, rather than a pangolin virus, remains the most probable origin of SARS-CoV-2 (Lau et al., 2020). Given the evolutionary gap, raTG13 is unlikely to be the direct ancestor virus of SARS-CoV-2, rather it branched out from the same ancestor (Xiaolu Tang, 2020; Zhou et al., 2020a). To compare the usage of ACE2 from different species by SARS-CoV-2 and raTG13, we transfected 293T cells with ACE2 of 10 different species and then infected them with pseudoviruses bearing SARS-CoV-2 S, SARS-CoV-2 S ΔPRRA, raTG13 S, raTG13 S+PRRA, and Pangolin GX S. Bald virus (envelope free) and pseudovirus bearing the vesicular stomatitis virus glycoprotein G (VSV-Gpp) were included in the study as negative and positive control. Pseudovirions bearing SARS-CoV-2 spike efficiently infected all but mouse ACE2 expressing cells. The same observation was made with pseudoviruses bearing SARS-CoV-2 S ΔPRRA and raTG13 S. Most strikingly, pseudovirus bearing raTG13 S+PRRA lost the ability to infect cells via pangolin and horseshoe bat ACE2 but gained infectivity via mouse ACE2 (Fig. 4). Pseudovirions bearing pangolin Spike efficiently entered cells via all ACE2 (including mouse ACE2 albeit by a one-log10 reduction). The above experiment was repeated in the presence of TMPRSS2 and same observations were made (Fig. S3).

**Figure 4.**
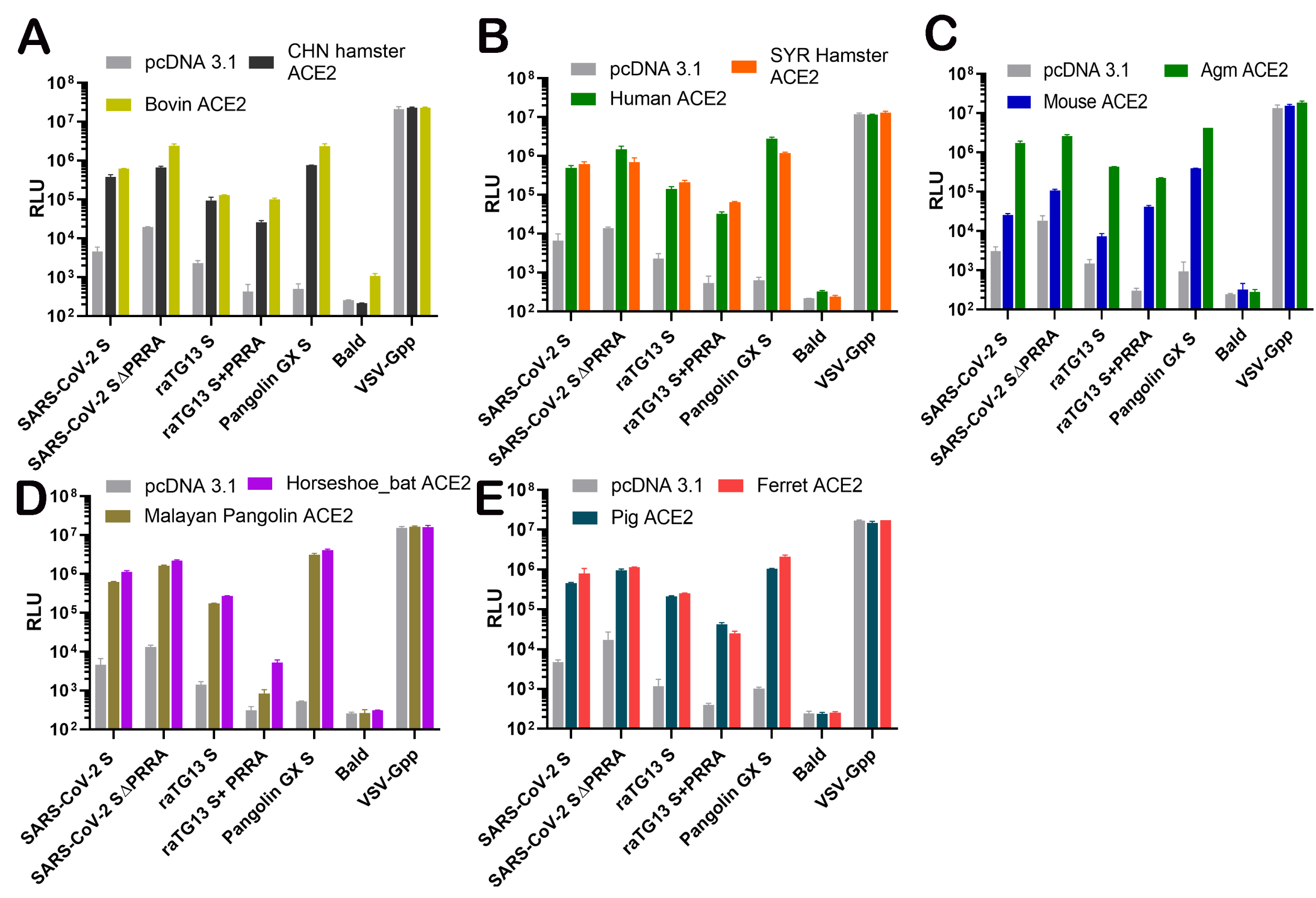
The novel S1/S2 site alters the dependence of raTG13 pseudovirus on ACE2 orthologs. Infectivity of pseudovirons harboring the indicated viral spike proteins on 293T cells transiently transfected with Chinese hamster ACE2, Bovine ACE2 (A); Syrian hamster ACE2, Human ACE2 (B); Agm ACE2, Mouse ACE2 (C); Horseshoe bat ACE2, Malayan Pangolin ACE2 (D); and Ferret ACE2, Pig ACE2 (E). Data are represented as mean ± SEM.

## Discussion

Here we experimentally demonstrated that SARS-CoV-2 is capable of infecting cells via ACE2 of a diverse range of species. Noticeably, a recent bioinformatics analysis ranked horseshoe bat and ferret ACE2 in the low or very low group in terms of the likelihood of bind SARS-CoV-2 (Damas et al., 2020), yet we found that SARS-COV-2 efficiently utilized both ACE2 to enter cells. Such a sharp contrast highlights the unpredictability of animal ACE2 that could be used by SARS-CoV-2 to enter cells based on current structural or sequence information. A secondary finding is that besides SARS-CoV-2, pseudoviruses bearing both bat raTG13 S and Pangolin GX S entered cells via a wide range of ACE2. Pangolin GX spike seems to use ACE2 from an even broader range of species to gain entry, at least in the case of mouse ACE2. Interestingly, despite a 87% identity to SARS-CoV-2 in the RBD region (Shang et al., 2020b), the RBM of pangolin/GX spike protein contains both residues that are favorable for ACE2 recognition (Leu455 and the 482–485 loop) and those that are less favorable for ACE2 recognition (Leu486, Glu493 and Thr501). Therefore, it was predicted that CoV-pangolin/GX may not recognize human ACE2 (Shang et al., 2020b). Yet our results here showed otherwise. Overall, our findings support potential cross-species transmissions of bat or pangolin coronaviruses, but also highlight the importance of surveying animal coronaviruses for their abilities to infect human cells.

This study also revealed a rather convoluted role of the PRRA insert between S1 and S2 subunits in SARS-CoV-2 entry. The presence of PRRA in SARS-CoV-2 clearly resulted in one (or two) cleavage site(s). However, at least on 293T-hACE2 cells, pseudovirus bearing SARS-CoV-2 S ΔPRRA exhibited even greater infectivity, suggesting that PRRA is dispensable to the Spike-driven viral entry. The raTG13 spike, which does not contain the PRRA sequence, is readily cleavable under the same condition. Addition of PRRA to raTG13 Spike further enhanced the cleavage but decreased the relevant pseudovirus infectivity on 293T-hACE2 cells, indicating that addition of PRRA does not enhance raTG13 Spike-driven entry. It is worth mentioning that SARS-CoV-2 and raTG13 spike protein sequences are identical within 76 residues downstream and upstream of the PRRA S1/S2 site (Supplemental Information). We envision that SARS-CoV-2 and raTG13 spike may adopt different structure conformations near the S1/S2 boundary such that the exact region within SARS-CoV-2 is less accessible as opposed to that in raTG13. Nonetheless, the PRRA insert and its mediated cleavage is clearly dispensable for pseudovirus to enter 293T-hACE2 cells. Having said that, SARS-CoV-2 spike protein does exhibit a basal level of fusogenic activity in the absence of ACE2, which is not seen with other spike proteins that were tested in this study. In consistence, we observed a basal level of infection of 293T cells by SARS-CoV-2 in the absence of exogenous ACE2 expression. Whether this reflects a receptor-independent route of entry or not needs to be further investigated, but it is possible that this basal level of fusogenic activity of SARS-CoV-2 could lead to cell-cell transmission. Overall, the spike protein-mediated cell-cell fusion in hACE2-293T cells largely correlates with the pseudovirus infectivity in the same cell type.

The PRRA insert appears to be essential for efficient Spike-dependent pseudovirus entry into the human lung epithelial cell line Calu-3. Two recent studies also reported similar findings (Hoffmann et al., 2020; Shang et al., 2020a). Such a sharp contrast with what was observed on 293T-hACE2 cells may be partially attributed to the relative abundance of proteases in cells. The acquisition of the 4 amino acid insert (PRRA) has been thought to broaden the activating protease repertoire of the SARS-CoV-2 S1/S2 cleavage site. Indeed, Jaimes et a.l. reported that the S1/S2 site of SARS-CoV-2 S is efficiently cleaved by furin-like, trypsin-like, and cathepsin proteases (Jaimes et al., 2020b). Hence, depending on the availability of the activating protease in each cell type, PRRA-guided cleavage may be subjected to differential regulations. In support of this notion, we noticed a partial cleavage of SARS-CoV-2 spike in 293T cells, whereas SARS-CoV-2 infected Caco-2 cells also exhibited much more extensive cleavage of spike although we cannot confirm whether this is mediated by the PRRA site. Astoundingly, similar cleavage occurs at a level that is barely detectible in Calu-3 cells despite it supports SARS-CoV-2 infection. RNA-seq analyses of SARS-CoV-2 infected Calu-3 and Caco-2 cells revealed different protease repertoires with cathepsin A being expressed to a much higher level in Caco-2 (Figure S4). It is conceivable that the spike protein is cleaved by different host proteases among 293T, Calu-3 and Caco-2. Given the very low level of cleavage that was observed in Calu-3 cells, it is plausible that the PRRA site becomes more important for the activating protease to cleave and prime Spike. Notably, CoVs enter the host cells via the endocytic pathway and non-endosomal pathway (Zumla et al., 2016). It is unclear which route is utilized for SARS-CoV-2 to enter lung cells, but the protease repertoires in endosomes versus that on the plasma membrane are expected to be different, which may further impact the infectivity of different pseudoviruses.

The most striking finding of this study is that insertion of PRRA into the raTG13 spike protein altered the ability of relevant pseudovirus to utilize ACE2 of three species to gain entry. Specifically, pseudovirus bearing raTG13 S+PRRA lost its ability to infect cells via pangolin and horseshoe bat ACE2 yet gained infectivity via mouse ACE2. The latter finding is particularly astounding given entry of all other pseudoviruses was somewhat impaired via mouse ACE2. The implications of this finding are twofold: First, if SARS-CoV-2 and raTG13 share the same ancestor which originates from horseshoe bat, it is likely that acquisition of PRAA would render this bat ancestor virus less efficient infecting horseshoe bat, hence the virus would have to find a new host. Secondly, insertion of PRRA may have a previously unrecognized impact on Spike-ACE2 interaction. Up to date, the interaction interface between SARS-CoV-2 spike protein and ACE2 lies between the receptor binding motif of spike protein and several residues of ACE2. Based on the known structure of SARS-CoV-2 spike protein, PRRA is part of a protruding loop that is rather distal to the RBD, hence is not expected to modulate Spike-ACE2 interaction. Binding of RBD to ACE2 locks the spike protein into a conformation such that the S1/S2 becomes more accessible to proteolytic cleavage and hence primes the fusion peptide for subsequent viral-cell fusion. Modeling SARS-CoV-2 Spike and mouse ACE2 interaction predicts that mouse ACE2 is unlikely to support entry, which has been widely verified in experiments including this one (Wan et al., 2020; Zhou et al., 2020b). Our findings, however, suggest raTG13 Spike may adopt a different conformation from SARS-CoV-2 Spike and the presence of PRRA may subtly modulate the binding of its RBD to ACE2 of horseshoe bat, pangolin and mouse.

In summary, we showed that spike proteins from all three viruses, SARS-CoV-2, bat CoV raTG13, and CoV-pangolin/GX, have the potential to mediate entry using ACE2 from multiple animal species besides human. The PRRA insertion selectively allows SARS-CoV-2 to infect human lung cell line Calu-3 and unexpected altered dependence of raTG13 Spike on ACE2 of three species.

## Acknowledgements

The following reagent was deposited by the Centers for Diseases Control and Prevention and obtained through BEI Resources, NIAID, NIH: SARS-Related Coronavirus 2, Isolate USA-WA1/2020, NR-52281. We are grateful to Dr. P. Shi (UTMB) for the generous gift of the mNeonGreen SARS-CoV-2.

## Author Contributions

S. L., P. S., C. L., W. W., C.C., and T.W. conducted the experiments; S. L., P. S. and T.W. analyzed the data, T. W. designed the experiments and wrote the paper.

## Figure Legends

**Figure S1. Expression of ACE2 orthologs and infection of Calu-3 and Caco-2**, Related to Figure 1 and Figure 2. (A) The expression of ACE2 ortholog plasmids in 293T cells were detected using mouse anti-V5 tag monoclonal antibody targeting the C-terminal V5 tag (in Red). Green, anti-β-actin antibody. (B) Representative confocal images of Huh7.5.1, Caco-2 and Calu-3 cells infected with icSARS-CoV-2-mNG at multiplicity of infection of 1 for 48 h.

**Figure S2**. (A) 293T cells or 293T-hACE2 cells were infected by pseudovirus bearing SARS-CoV-2 S. Bald virus without any viral envelope or VSV-Gpp were included as negative and positive control. (B) Entry of MLV pseudotyped with SARS-CoV-2 S, SARS-CoV-2 S ΔPRRA in Vero cells.

**Figure S3. The novel S1/S2 site alters the dependence of raTG13 pseudovirus on ACE2 orthologs**, related to Figure 4. 293T cells expressing each ACE2 ortholog with TMPRSS2 were infected by pseudoviruses harboring SARS-CoV-2 S (A), SARS-CoV-2 S ΔPRRA (B), raTG13 S (C), raTG13 S+PRRA (D), pangolin S (E), VSV-G (F). Luciferase activities in cell lysates were determined at 48 h post infection. Data are represented as mean ± SEM.

**Figure S4**. Protease repertoires in Calu-3 and Caco-2. Calu-3 and Caco-2 cells were infected with SARS-CoV-2 USA/WA1-2020 at MOI of 0.1 for 4, 24, and 48 hours. RNA was isolated from uninfected and infected cells were subject to mRNA-Seq. Reads per kilobase per million mapped reads (RPKM) was calculated to show expression level of proteases that may cleave SARS-CoV-2 spike protein at the S1/S2 site. Also refer to Table S1.

## STAR Methods

### Resource Availability

#### Lead Contact

Further information and requests for reagents may be directed to and will be fulfilled by Lead Contact Tony Wang.

#### Materials Availability

All unique/stable reagents generated in this study are available from the Lead Contact with a completed Materials Transfer Agreement.

#### Data and Code Availability

The RNA-seq datasets of SARS-CoV-2 USA/WA1-2020 infected Calu-3 and Caco-2 cells have been submitted as Table S1.

### Experimental Model and Subject Details

#### Viruses and Cells

Vero E6 cell line (Cat # CRL-1586) and Caco-2 (Cat # HTB-37) cell line were purchased from American Type and Cell Collection (ATCC) and cultured in eagle’s minimal essential medium (MEM) supplemented with 10% fetal bovine serum (Invitrogen) and 1% penicillin/streptomycin and L-glutamine. Calu-3 cell line (Cat # HTB-55) was obtained from ATCC and maintained in EMEM+20%FBS. Huh7.5.1 was a gift of F. Chisari and maintained in (DMEM+10% FBS) (Zhong et al., 2005).

The SARS-CoV-2 isolate USA-WA1/2020 was obtained from BEI Resources, NIAID, NIH, and had been passed three times on Vero cells and 1 time on Vero E6 cells prior to acquisition. It was further passed once on Vero E6 cells in our lab. The virus has been sequenced verified to contain no mutation to its original seed virus. A reporter virus inserting the monomeric neon green (mNG) in the SARS-CoV-2 USA-WA1/2020 backbone was obtained from Dr. Pei-Yong Shi’s laboratory (university of Texas Medical Branch) at passage 2 and was further expanded once on Vero E6 (Xie et al., 2020) at FDA.

#### Virus titration

SARS-CoV-2 isolate USA-WA1/2020 was titered using the Reed & Muench Tissue Culture Infectious Dose 50 Assay (TCID50/ml) system (L.J. Reed, 1938). Vero cells were plated the day before infection into 96 well plates at 1.5 × 104 cells/well. On the day of the experiment, serial dilutions of virus were made in media and a total of 6-8 wells were infected with each serial dilution of the virus. After 48 hours incubation, cells were fixed in 4% paraformaldehyde followed by staining with 0.1% crystal violet. The TCID50 was then calculated using the formula: log (TCID50) = log(do)+ log (R)(f+1). Where do represents the dilution giving a positive well, f is a number derived from the number of positive wells calculated by a moving average, and R is the dilution factor.

#### Constructs

Full-length SARS-CoV-2 spike (YP_009724390.1), Bat raTG13 Spike (QHR63300.2), Pangolin Guangxi spike (QIA48614) were human codon-optimized and synthesized by GenScript. Human ACE2 (NM_021804.2) was purchased from GenScript with a C-terminal flag tag. Other ACE2 genes were synthesized (GenScript Biotech) and subcloned into the pcDNA3.1(+) vector between BamHI and XhoI sites with a C-terminal V5 tag. Sequences of all spike proteins and ACE2 orthologs can be found in Supplemental Information. TMPRSS2, psPAX2, pCMV-CREM and pSV-STOP-luc were purchased from Addgene.

SARS-CoV-2 S ΔPRRA was generated using overlap PCR and cloned into pcDNA3.1(+) vector between BamHI and XhoI sites. The sequences of the primers are: SARS-2S-F 5’-GCTCGGATCCCCACCATGTT-3’, SARS-2S-PRRA R: 5’-CTTGCCACAGACCGGGAGTTTGTCTGGGTCTGGTA-3’, SARS-2S-PRRA F: 5’-AGACAAACTCCCGGTCTGTGGCAAGCCAGTC-3’, SARS-2S-R: 5’-TCTAGACTCGAGTTAGGTGT3’. The raTG13 S+PRRA mutation was created by overlap PCR. Mutant PCR fragment was digested by BamHI/XhoI and ligated into BamHI/XhoI digested pcDNA3.1 vector. The sequences of the primers are as follows:

RaTG13 S F: 5’-GCGGATCCCCACCATGTTCGTGTTTCTGGTGCT-3’

RaTG13 S PRRA R: 5’-GGCCACGCTCCTTGCTCTCCTTGGAGAGTTTGTCTGGGTCTGAT3’

RaTG13 S PRRA F: 5’-CAAACTCTCCAAGGAGAGCAAGGAGCGTGGCCTCCCAGTC-3’

RaTG13 S R: 5’-ATATAGCTCGAGTTAGGTATAGTGCAG-3’

#### SARS-CoV-2 pseudovirus production

Human codon-optimized cDNA encoding SARS-CoV-2 S glycoprotein (NC_045512) was synthesized by GenScript and cloned into eukaryotic cell expression vector pcDNA 3.1 between the BamHI and XhoI sites. Pseudovirions were produced by co-transfection Lenti-X 293T cells with psPAX2, pTRIP-luc, and SARS-CoV-2 S expressing plasmid using Lipofectamine 3000. The supernatants were harvested at 48h and 72h post transfection and filtered through 0.45-mm membranes. To pellet down pseudovirions, the viral like particles were centrifuged at 20,000 xg for 2 h at 4 °C, then resuspend in SDS-PAGE sample buffer after removing all supernatant. Alternatively, pseudovirions were produced by co-transfection Lenti-X 293T cells with pMLV-gag-pol, pFBluc, and pcDNA 3.1 SARS-CoV-2 S using Lipofectamine 3000. The supernatants were harvested at 48h and 72h post transfection and filtered through 0.45-mm membranes. The MLV-based pseudovirus infects VeroE6 cells more efficiently than HIV-based counterpart.

#### CoV Spike-Mediated Cell–Cell Fusion Assay

In brief, 293T cells were transfected with hACE2 and Stop-Luc construct which contains a firefly luciferase reporter gene whose transcription is prevented by a Stop cassette flanked by LoxP sites. 24 h post-transfection, these recipient cells were mixed at a 1:1 ratio with 293T cells expressing Cre and CoV S (donor cells) to initiate cell–cell fusion. Luciferase activity was measured 48 h thereafter.

#### Immunofluorescence assay

Cells were plated on collagen-coated glass coverslips the day before infection into 24 well plates at 5 × 104 cells/well. The cells were washed for 5 minutes 3 times with 1X phosphate buffered saline. After wash, cells were fixed in 4% paraformaldehyde for 15 minutes at room temperature. Images were captured by a Zeiss LSM 880 laser scanning confocal microscope.

#### Flow cytometry

Infected cells were trypsinized and washes in 1x PBS. mNG+ cells were quantified by BD FACSCanto II (BD Biosciences). Data was analyzed by FlowJo software.

#### Immunoblotting

Cell lysates were prepared with RIPA buffer (50 mM Tris-HCl [pH 7.4]; 1% NP-40; 0.25% sodium deoxycholate; 150 mM NaCl; 1 mM EDTA; protease inhibitor cocktail (Sigma); 1 mM sodium orthovanadate), and insoluble material was precipitated by brief centrifugation. Protein concentration of lysates was determined by BCA protein assay (Thermo Scientific). Lysates containing equal amounts of protein were loaded onto 4-20% SDS-PAGE gels and transferred to a nitrocellulose membrane (LI-COR, Lincoln, NE), blocked with 10% milk for 1 h, and incubated with the primary antibody overnight at 4 °C. Membranes were blocked with Odyssey Blocking buffer (LI-COR, Lincoln, NE), followed by incubation with primary antibodies at 1:1000 dilutions. Membranes were washed three times with 1X PBS containing 0.05% Tween20 (v/v), incubated with IRDye secondary antibodies (LI-COR, Lincoln, NE) for 1 h, and washed again to remove unbound antibody. Images were captured by the Odyssey CLx infrared imaging system (LI-COR Biosystems, Lincoln, NE).

#### mRNA-Seq by the next generation sequencing

Total RNA extracted from Calu-3 and Caco-2 cells were used for sequencing library preparation. RNA quality was assessed by Agilent 2100 Bioanalyzer (Agilent Technologies, Santa Clara, CA, USA) and the RNA integration numbers (RIN) were all greater than 9. Poly-A enriched stranded libraries were generated using TruSeq Stranded mRNA Library Preparation kit (Illumina, San Diego, CA, USA), following Illumina’s protocol. In brief, 0.5 µg of total RNA was poly-A enriched, fragmented, and reverse transcribed into cDNAs. Double stranded cDNAs were adenylated at the 3′ ends and individually indexed followed by 15-cycle PCR amplification. The quality and yield of prepared libraries were evaluated using Agilent 2100 Bioanalyzer and Qubit 4 (Thermo Fisher Scientific). Libraries of mRNA-Seq were sequenced on an Illumina NovaSeq 6000 instrument following the manufacture’s instruction. Data analysis was performed using RNA-Seq tool of CLC Genomics Workbench V12 (QIAGEN, Hilden, Germany), which aligned the paired-end reads to the human hg38 reference genome and calculated the gene expression. For each sample, reads per kilobase per million mapped reads (RPKM) was calculated to show expression level. The full dataset can be found in Table S1.

#### Statistical analysis

All experiments were independently performed at least twice times as indicated in the Figure legends. Except were specified, bar graphs were plotted to show mean ± standard deviation (SD). Statistical analyses were performed using Prism 8.

---

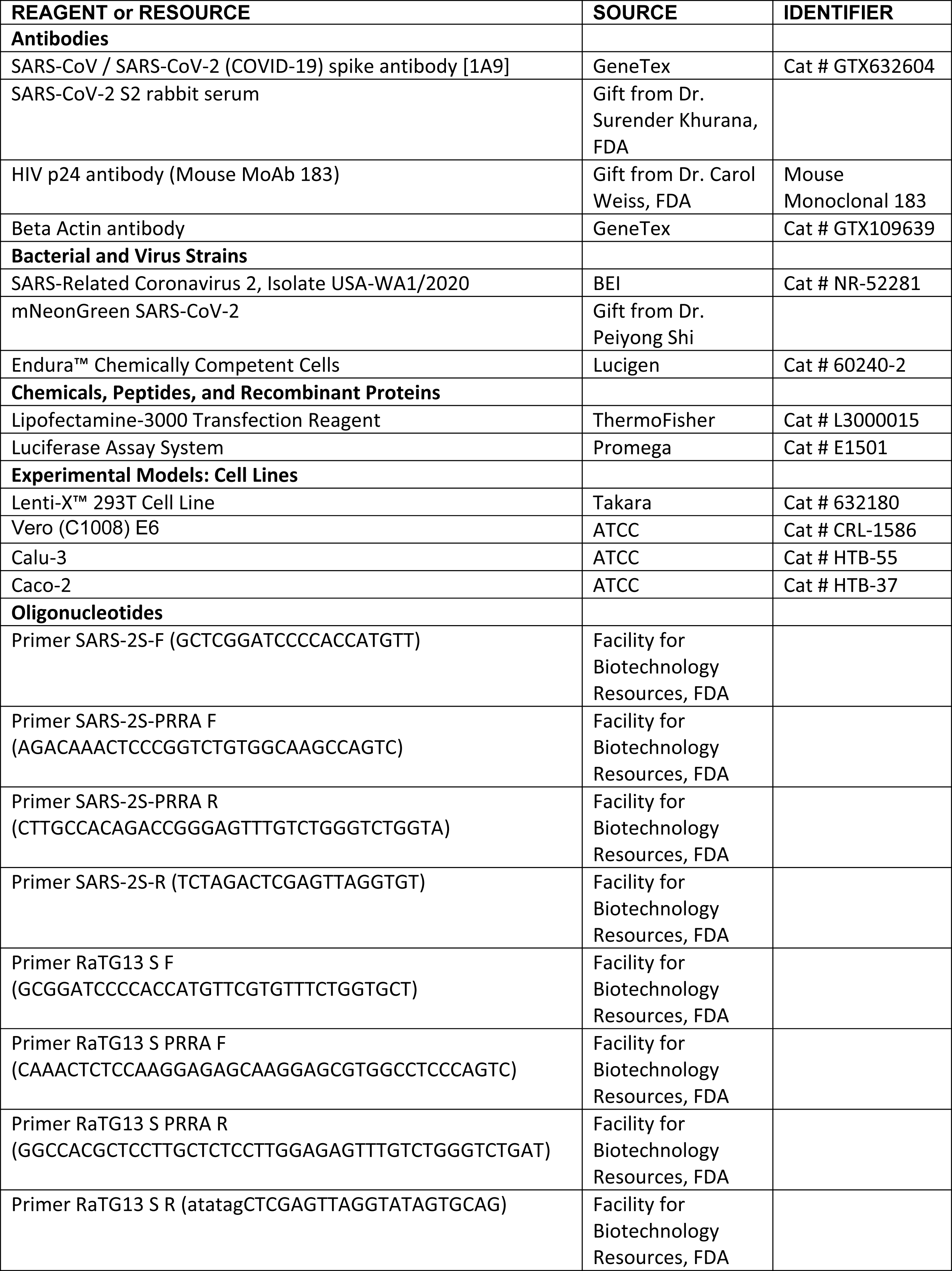

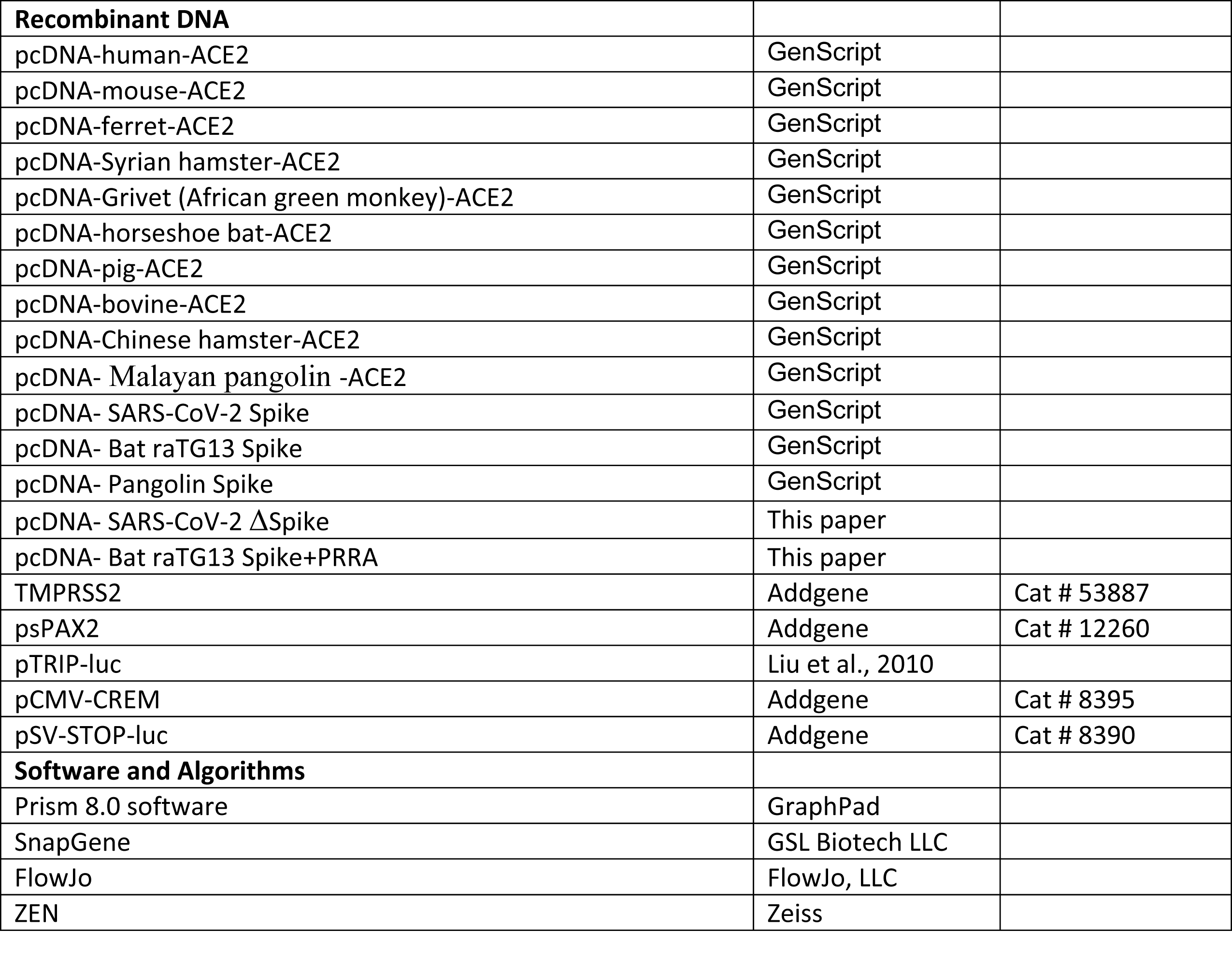

